# Dietary cardenolides enhance growth and change the direction of the fecundity-longevity trade-off in milkweed bugs (Heteroptera: Lygaeinae)

**DOI:** 10.1101/2021.03.29.437508

**Authors:** Prayan Pokharel, Anke Steppuhn, Georg Petschenka

**Affiliations:** Department of Applied Entomology, Institute for Phytomedicine, University of Hohenheim, Otto-Sander-Str. 5, 70599 Stuttgart, Germany; Department of Molecular Botany, Institute for Biology, University of Hohenheim, Garbenstr. 30, 70599 Stuttgart, Germany

**Keywords:** Cardenolides, Fitness costs, Life-history traits, Milkweed bugs, Na^+^/K^+^-ATPase, Sequestration, Trade-off

## Abstract

1. Sequestration, i.e., the accumulation of plant toxins into body tissues for defence, is primarily observed in specialised insects. Sequestration was frequently predicted to incur a physiological cost mediated by increased exposure to plant toxins and may require resistance traits different from those of non-sequestering insects. Alternatively, sequestering species could experience a cost in the absence of toxins due to selection on physiological homeostasis under permanent exposure of sequestered toxins in body tissues.
2. Milkweed bugs (Heteroptera: Lygaeinae) sequester high amounts of plant-derived cardenolides. Although being potent inhibitors of the ubiquitous animal enzyme Na^+^/K^+^-ATPase, milkweed bugs can tolerate cardenolides by means of resistant Na^+^/K^+^-ATPases. Both adaptations, resistance and sequestration, are ancestral traits shared by most species of the Lygaeinae.
3. Using four milkweed bug species and the related European firebug (*Pyrrhocoris apterus*) showing different combinations of the traits ‘cardenolide resistance’ and ‘cardenolide sequestration’, we set out to test how the two traits affect larval growth upon exposure to dietary cardenolides in an artificial diet system. While cardenolides impaired the growth of *P. apterus* nymphs neither possessing a resistant Na^+^/K^+^-ATPase nor sequestering cardenolides, growth was not affected in the non-sequestering milkweed bug *Arocatus longiceps*, which possesses a resistant Na^+^/K^+^-ATPase. Remarkably, cardenolides increased growth in the sequestering dietary specialists *Caenocoris nerii* and *Oncopeltus fasciatus* but not in the sequestering dietary generalist *Spilostethus pandurus*, which all possess a resistant Na^+^/K^+^-ATPase.
4. We then assessed the effect of dietary cardenolides on additional life history parameters, including developmental speed, the longevity of adults, and reproductive success in *O. fasciatus*. Remarkably, nymphs under cardenolide exposure developed substantially faster and lived longer as adults. However, fecundity of adults was reduced when maintained on cardenolide-containing diet for their entire life-time but not when adults were transferred to non-toxic sunflower seeds.
5. We speculate that the resistant Na^+^/K^+^-ATPase of milkweed bugs is selected for working optimally in a ‘toxic environment’, i.e. when sequestered cardenolides are stored in the body tissues. Contrary to the assumption that toxins sequestered for defence produce a physiological burden, our data suggest that they can even increase fitness in specialised insects.

## 1. Introduction

Chemical defences are widespread among animals and remarkably diverse across species. Many insect herbivores accumulate secondary metabolites from their host plants and utilize them for their defence to ward off natural enemies, a phenomenon called sequestration (Petschenka and Agrawal 2016). For example, caterpillars of the monarch butterfly (*Danaus plexippus*) sequester cardenolides from their host plant milkweed (*Asclepias* spp.). On the contrary, other insects produce their toxins via *de novo* synthesis as observed in leaf beetles (Chrysomelidae) (Pasteels, Braekman, and Daloze 1988) and cyanogenic butterflies (Heliconiinae) (Brown and Francini 1990). Sequestration of toxins from plants and *de novo* synthesis of toxins are negatively correlated traits in some insects (Engler-Chaouat and Gilbert 2007). In the light of evolutionary theory, these patterns are assumed to represent trade-offs between gained protection against predators and costs of possessing these defences (Bowers 1992; Camara 1997).

It was speculated that physiological costs of *de novo* synthesis of defensive compounds are higher compared to the costs of sequestration of plant toxins (Fürstenberg-Hägg et al. 2014), which may explain why sequestration is a common phenomenon currently reported for more than 250 insect species acquiring toxins from at least 40 plant families (Opitz and Müller 2009). Nevertheless, sequestration of chemical defences may incur physiological costs (Camara 1997; Reudler et al. 2015) since sequestering insects are exposed to high concentrations of toxins stored within their body tissues. However, empirical evidence on the benefits of defences is more apparent than their costs, and the costs of chemical defences are not always easy to detect (Ruxton 2014; Lindstedt et al. 2010). In line with this, evidence for actual physiological costs such as effects on growth or other fitness parameters like longevity and fecundity is very scarce (Zvereva and Kozlov 2016). Understanding the evolution of defences in insects will require rigorous comparative analyses of the specific adaptations underlying chemical defences.

Trade-offs play a crucial role in an organism’s life-history and occur when a beneficial change in one trait is linked to an unfavourable change in another trait with a possible cost (Stearns 1989). Life-history theory suggests that fitness determining traits such as longevity and fecundity are negatively associated with each other (Holliday 1994; Flatt 2011). However, results are contradictory with studies either showing positive, negative, or zero correlation between these two traits among individuals within a population (G Bell and Koufopanou 1986; Van Noordwijk and de Jong 1986). Generally, life-history trade-offs result from compromises in resource allocation across growth, survival, maintenance, and reproduction under challenges such as predation occurring in an ecosystem (Levins 1968; Sibly and Calow 1986; Roff 1992; Walsh and Reznick 2010). Regarding chemical defence, an organism’s potential physiological costs may be estimated as trade-offs between investments in defence and other physiological parameters such as growth, longevity, or fecundity (Camara 1997; Ruxton et al. 2019).

Cardenolides are produced by more than ten different plant families (Malcolm 1991; Luckner and Wichtl 2000) and are toxic to animals because they specifically inhibit the ubiquitous enzyme Na^+^/K^+^-ATPase (Lingrel 1992; Emery et al. 1998). Na^+^/K^+^-ATPase is a cation carrier responsible for essential physiological functions such as the generation of neuronal action potentials and maintenance of an electrochemical gradient across the cell membrane (Jorgensen, Håkansson, and Karlish 2003). Remarkably, insects from at least six orders, including milkweed bugs (Heteroptera: Lygaeinae), milkweed butterflies (Lepidoptera: Danaini), and leaf beetles (Coleoptera: Chrysomelidae), show common adaptations to cardenolides (Dobler et al. 2015). Resistance in these groups is mediated by target site insensitivity due to a few amino acid substitutions in the first extracellular loop of the alpha subunit of Na^+^/K^+^-ATPase (ATPα), resulting in a high level of molecular convergence, i.e., often the identical amino acid substitutions at the same positions confer resistance (Zhen et al. 2012; Dobler et al. 2012, 2015).

Milkweed bugs are seed feeders primarily found on Apocynaceae species and on unrelated cardenolide-producing plants (Petschenka et al. 2020). Besides feeding on cardenolide-containing plants, milkweed bugs also sequester cardenolides to ward off predators (Evans, Castoriades, and Badruddine 1986; Pokharel et al. 2020). Alteration of the Na^+^/K^+^-ATPase in the Lygaeinae is probably correlated with several duplications of the ATPα1 gene resulting in four ATPα1 paralogs (A, B, C, D) found in *Oncopeltus fasciatus* and *Lygaeus kalmii* (Yang et al. 2019) (Zhen et al. 2012; Dalla and Dobler 2016). Cardenolide-resistant Na^+^/K^+^-ATPases and the ability to sequester cardenolides are most likely synapomorphic features in the Lygaeinae, which may account for the milkweed bugs’ evolutionary success (Bramer et al. 2015).

The goal of our study was to evaluate if cardenolide exposure and sequestration causes physiological costs or benefits, and if these effects differ across closely related milkweed bug species possessing different combinations of the traits ‘resistance’ and ‘sequestration’ (i.e., having resistant/sensitive Na^+^/K^+^-ATPases and sequestering/not sequestering). Our set of species included dietary specialist and generalist milkweed bug species and the European linden bug *Pyrrhocoris apterus* (Linnaeus, 1758, Pyrrhocoride) having no adaptations to cardenolides for comparison (Figure 1).

**Figure 1.**
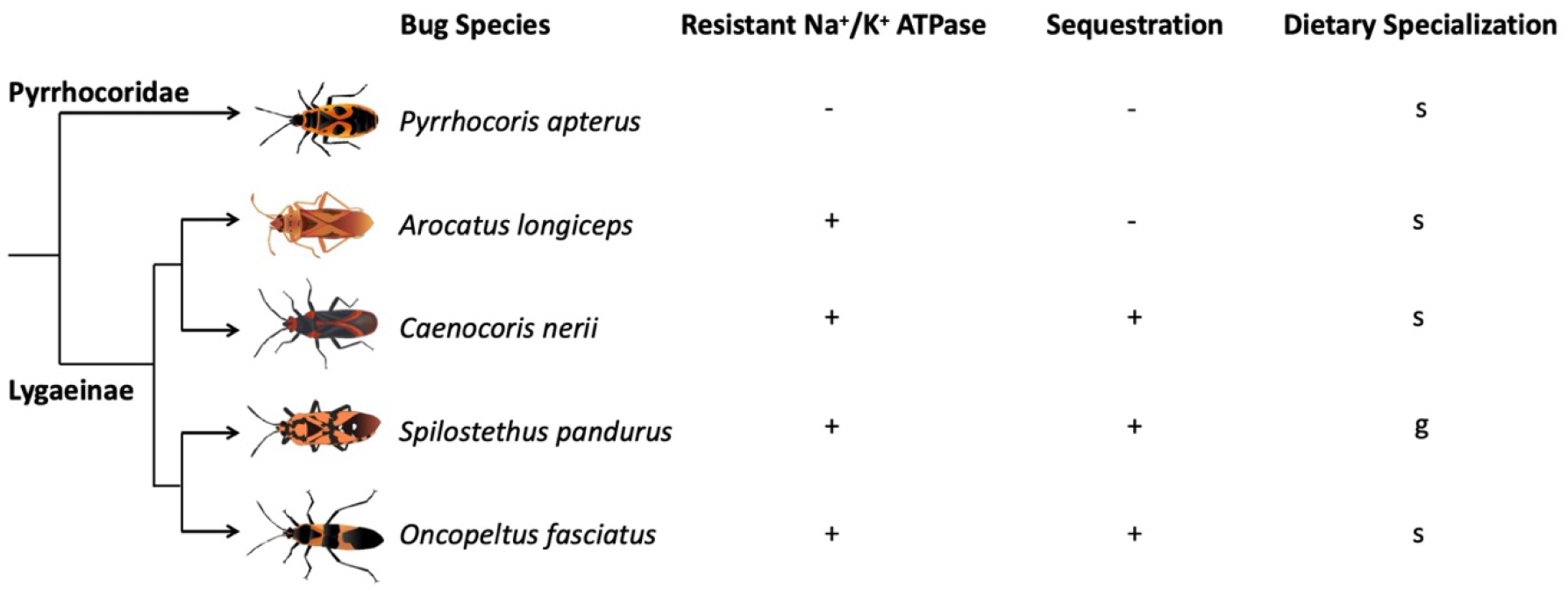
Overview of the Heteroptera species used in the experiments with their key traits. We compared four species of Lygaeinae sharing cardenolide resistant Na^+^/K^+^-ATPase (+ = resistant; - = sensitive) but differing in their ability to sequester cardenolides (+ = sequestering; - = not sequestering). Within the sequestering milkweed bug species, *O. fasciatus* and *C. nerii* may be classified as host-plant specialists (s = specialist), while *S. pandurus* uses a wide variety of host plants (g = generalist). *A. longiceps* is specialised on *Platanus* and *Ulmus* which are devoid of cardenolides, and lost its ability to sequester cardenolides in the course of evolution. *Pyrrhocoris apterus* belongs to the relatively closely related family Pyrrhocoridae and is specialised on cardenolide-free Malvaceae. Furthermore, it is known to possess a sensitive Na^+^/K^+^-ATPase, and, based on our recent analyses, does not sequester cardenolides. Phylogenetic relationships of Heteroptera species are based on Bramer et al. 2015.

The milkweed bug species we used were *O. fasciatus* (Dallas, 1852), *Caenocoris nerii* (Germar, 1847) (both resistant and sequestering, dietary specialists), *Spilotethus pandurus* (Scopoli, 1763) (resistant and sequestering, dietary generalist), and *Arocatus longiceps* (Stal, 1872) (resistant and not sequestering, dietary specialist). As Stearns (1992) considers manipulating a single factor the most reliable and informative method to measure costs (or trade-offs, if any), we manipulated cardenolide concentration in the diet as a single factor. For this purpose, we established an artificial diet approach and raised larvae on increasing doses of cardenolides in the diet. We assessed growth over the course of development. Furthermore, we determined the amount of sequestered cardenolides using liquid chromatography (HPLC-DAD).

To test for effects on potential trade-offs between life-history parameters due to dietary cardenolides, we investigated the influence of dietary cardenolides on the longevity and fecundity in *O. fasciatus*. Trade-offs have been measured in the field (Clutton-Brock 1982; Clutton-Brock, Guinness, and Albon 1983) and in the laboratory (Partridge and Farquhar 1981), e.g. by genotypic studies in *Drosophila melanogaster* (Rose and Charlesworth 1981a, 1981b), and by phenotypic studies in *Daphnia pulex* and *Platyias patulus* (Graham Bell 1984a, 1984b). How different traits will trade-off depends on ecological factors and the physiological state of an organism. Thus, trade-offs can change across different environments in different species or even within the same species. Therefore, a life-history trade-off probably only appears in a species under a particular set of conditions, such as stress (Reznick 1985).

To explicitly investigate the influence of dietary toxins in milkweed bug species, we set out to test the following hypotheses: i) insect species having different physiological traits (resistance and sequestration) and ecological strategies (generalist and specialist) will react differently to dietary toxins, ii) sequestering species will experience costs, either in the presence or absence of toxins, and iii) the fecundity-longevity trade-off will be altered by dietary toxins.

## 2. Materials and Methods

### 2.1. Preparation of artificial diet

We followed Jones et al.’s method to prepare an artificial diet for *Oncopeltus fasciatus* (Jones, Mel, and Yin 1986) but used a modified approach to offer the diet to the bugs. Sunflower seeds (25 g), wheat germ (25 g), casein (25 g), sucrose (10 g), Wesson’s salt (4 g), vitamins (Vanderzant Vitamin mix, 5 g), methyl 4-hydroxybenzoate (1 g), sorbic acid (0.5 g), olive oil (7.5 g), and toxins (only for the treatment groups, not for controls) were blended in 200 ml of water until the mixture was homogenous. Agar (7.5 g) was boiled separately in 300 ml of water in a microwave. After 5 min, when the agar mixture had slightly cooled down, the agar and the mixture were combined and blended again. The obtained paste was poured into plastic boxes and stored at 4 °C for further use. For the feeding assays, an aliquot of diet was filled into the lid of a 2 ml Eppendorf tube and sealed with a piece of stretched parafilm to be used as an ‘artificial seed’. Using portions from the same initial bulk of the diet, we added increasing amounts of an equimolar mixture of crystalline ouabain and digitoxin (Sigma-Aldrich, Taufkirchen, Germany) to prepare diets with either 2 (‘low’), 6 (‘medium’), or 10 (‘high’) mg cardenolide/mg dry weight of diet. The initial diet without cardenolides added was used as a control. We used the polar ouabain and the relatively apolar digitoxin to mimic the condition that plants typically produce an array of cardenolides with a wide polarity spectrum. The concentration of cardenolides in the diet was chosen to be in the range of natural cardenolide concentrations observed in *Asclepias* seeds (Isman 1977). For each batch of diet prepared, we verified the concentrations of cardenolides across all dietary treatments by using high-performance liquid chromatography (HPLC, see Section 2.2).

### 2.2. Quantification of cardenolides

To verify the amount of cardenolides in the artificial diet, cubes of diet (approx. 20-25 mg dry weight) were freeze-dried, weighed, and added to a 2 ml screw-capped tube containing approximately 900 mg of zirconia/silica beads (ø 2.3 mm, BioSpec Products, Inc., Bartlesville, OK US). One ml HPLC-grade methanol containing 0.01 mg/ml of oleandrin (PhytoLab GmbH & Co. KG, Vestenbergsgreuth, Germany) as an internal standard was added to the tube and diet samples were homogenized for two cycles of 45 s at 6.5 m/s in a Fast Prep™ homogenizer (MP Biomedicals, LLC, Solon, OH, US). Supernatants were transferred into fresh tubes after centrifugation at 16,100 g for 3 min. Original samples were extracted two more times with 1 ml of pure methanol as described above. All the supernatants of a sample were combined, evaporated to dryness under a stream of nitrogen gas, and resuspended with 100 μl methanol by agitating tubes in the Fast Prep™ homogenizer without beads. Subsequently, samples were filtered into HPLC vials using Rotilabo ® syringe filters (nylon, pore size: 0.45 μm, ø 13 mm, Carl Roth GmbH & Co. KG, Karlsruhe, Germany). Finally, 15 μl of the extract was injected into an Agilent 1100 series HPLC (Agilent Technologies, Santa Clara, US) equipped with a photodiode array detector, and compounds were separated on an EC 150/4.6 NUCLEODUR^®^ C18 Gravity column (3 μm, 150 mm × 4.6 mm, Macherey-Nagel, Düren, Germany). Cardenolides were eluted at a constant flow rate of 0.7 ml/min at 30°C using the following acetonitrile-water gradient: 0–2 min 10% acetonitrile, 13 min 95% acetonitrile, 18 min 95% acetonitrile, 23 min 10% acetonitrile, 5 min reconditioning at 10% acetonitrile. The same HPLC method was used for quantification of sequestered cardenolides in *O. fasciatus* and *P. apterus*, respectively. For the analysis of sequestered cardenolides in *C. nerii*, *S. pandurus* and *A. longiceps*, we used a different acetonitrile-water gradient to achieve improved separation of polar cardenolides than digitoxin: 0–2 min 16% acetonitrile, 25 min 70% acetonitrile, 30 min 95% acetonitrile, 35 min 95% acetonitrile, 37 min 16% acetonitrile, 10 min reconditioning at 16% acetonitrile. We interpreted peaks with symmetrical absorption maxima between 216 and 222 nm as cardenolides (Stephen B. Malcolm and Zalucki 1996) and integrated peaks at 218 nm using the Agilent ChemStation software (B.04.03). The amount of cardenolides in a sample was quantified based on the peak area of the known concentration of the internal standard oleandrin.

### 2.3. Insect colonies

*O.fasciatus* were obtained from a long-term laboratory colony (originally from the United States) maintained on sunflower seeds. We collected specimens of *P. apterus* in the vicinity of linden trees (*Tilia* spp., Malvaceae) and specimens of *A. longiceps* from under the bark of plane trees (*Platanus* spp., Platanaceae) in Giessen, Germany. Specimens of *S. pandurus* and *C. nerii* were collected from a *Nerium oleander* habitat close to Francavilla di Sicilia, Messina, Sicily, Italy. In the laboratory, insect colonies were reared in plastic boxes (19 × 19 × 19 cm) covered with gauze in a climate chamber (Fitotron® SGC 120, Weiss Technik, Loughborough, UK) at 27°C, 60% humidity and a day/night cycle of 16/8 h under artificial light. We reared all insects on organic sunflower seeds (Alnatura GmbH, Darmstadt, Germany), supplied water in cotton-plugged Eppendorf tubes and included a piece of cotton as a substrate for oviposition. In addition to sunflower seeds, *P. apterus* was provided with approximately 10 freshly chopped mealworms twice a week.

For the experiments described below, we used first-generation offspring from field-collected *P.apterus* and *A. longiceps* (maintained as described above), whereas *S. pandurus* and *C. nerii* offspring were obtained from colonies maintained in the lab for more than four generations.

### 2.4. Growth Assay

We carried out feeding assays to investigate the influence of increasing doses of dietary toxins on the growth of larvae of four species of milkweed bugs (*O. fasciatus*, *C. nerii, S. pandurus*, and *A. longiceps*) and an outgroup, *P. apterus*. These species either lack the ability to tolerate and sequester cardenolides (*P. apterus*), possess a cardenolide-resistant Na^+^/K^+^-ATPase, and can sequester cardenolides (*O. fasciatus, C. nerii* and *S. pandurus*) or possess a cardenolide-resistant Na^+^/K^+^-ATPase but lost the ability to sequester (*A. longiceps*) (Figure 1). We placed three 2^nd^ instar (L2) larvae from the stock colonies in a Petri-dish (60 mm × 15 mm, with vents, Greiner Bio-One, Frickenhausen, Germany) lined with filter paper (Rotilabo^®^-round filters, Carl Roth GmbH & Co. KG, Karlsruhe, Germany) that was supplied with an artificial seed (either being devoid of cardenolides, or possessing a cardenolide concentration of 2, 6, or 10 μg/mg dry weight) and a water source (a 0.5 ml Eppendorf tube plugged with cotton). The artificial seeds were replaced once in two weeks. All Petri-dishes were spatially randomized and maintained in a controlled environment (KBWF 240 climate chamber, Binder, Tuttlingen, Germany) at 21°C, 60% humidity and a day/night cycle of 16/8 h under artificial light. The growth of larvae was assessed twice a week over a period of three weeks by sedating all bugs of a Petri-dish with CO2 and weighing them jointly. After reaching adulthood, at least one bug per Petri-dish was transferred to a toxin-free diet for 10 days to avoid a potential bias from toxins remaining in the gut (i.e., not being sequestered) by purging. Finally, bugs were killed by freezing, freeze-dried, weighed, extracted, and analysed by HPLC to estimate the amount of sequestered cardenolides (see Section 2.2). We also estimated the amount of excretion products on filter paper that may provide an indication of food intake (see supplementary methods). For *O. fasciatus*, growth assays were performed in three batches, and estimation of sequestered cardenolides was performed in two batches.

### 2.5. Life-history assays with *O. fasciatus*

#### 2.5.1. Developmental time

Since the effects of dietary cardenolides on growth were most pronounced in *O. fasciatus*, we carried out a separate experiment to assess additional life-history parameters such as duration of larval development, adult lifespan and body size under the influence of dietary cardenolides in this species. We chose the medium-dose cardenolide (6 μg/mg dry weight) because we observed that *O. fasciatus* showed maximal growth on this diet in our previous experiment. The experimental setup was similar to that of the growth assay. However, here only one L2 larva was placed in each Petri-dish to monitor its development time. Petri-dishes lined with filter paper either containing medium-dose diet or control diet without toxins and a water source (see Section 2.4) were spatially randomized and kept in a climate chamber (Fitotron^®^ SGC 120, Weiss Technik, Loughborough, UK) at 27°C, 60% humidity and a day/night cycle of 16/8 h under artificial light. We checked the Petri-dishes every day for dead individuals, raised the bugs until adulthood and observed them until they died. We also measured the body length of adult males and females raised on the two different diets using a Vernier calliper.

#### 2.5.2. Reproductive fitness

We conducted two additional experiments to assess the effect of dietary cardenolides on reproductive fitness. We raised L2 larvae in bulk (around 50 individuals) until adulthood in plastic boxes (19 × 19 × 19 cm) covered with gauze either on two artificial seeds of medium-dose or control diet and a water source (four Eppendorf tubes of 2 ml plugged with cotton). Boxes were kept in a climate chamber (Fitotron® SGC 120, Weiss Technik, Loughborough, UK) at 27°C, 60% humidity and a day/night cycle of 16/8 h under artificial light. Artificial seeds were replaced once in two weeks. At least three (but not older than six) days old males and females from the same treatment were paired in Petri-dishes (9 cm × 1.5 cm, with vents, Greiner Bio-One, Frickenhausen, Germany) lined with filter paper and a water source (a 2 ml Eppendorf tube plugged with cotton). Additionally, we included a piece of cotton as a substrate for oviposition. Petri-dishes were spatially randomized and kept under the same conditions as described above. In a first experiment, adult bugs were supplied with the same type of artificial diet that they were raised upon (i.e., either control or medium-dose cardenolide artificial seeds). In nature, adults of *O. fasciatus* disperse after reaching adulthood and forage for other seeds besides *Asclepias* spp. (Feir 1974). Therefore, we carried out a second experiment under the same conditions as described above in which pairs of bugs were supplied with approx. 20 sunflower seeds instead of artificial seeds. Since a substantial portion of eggs in both experiments were unviable (possibly due to the use of an artificial diet), we counted only hatchlings and not the total number of eggs produced by each female over its entire lifespan.

### 2.6. Statistical Analysis

Statistical analyses were computed using JMP® Pro 15 statistical software (SAS Institute, Cary, NC, US). All data were 1og_10_-transformed to achieve homogeneity of variances and normality of residuals. For feeding experiments, we analysed sequential data on larval masses with Repeated Measures ANOVA followed by LSMeans Tukey HSD test to assess potential differences across treatments. We compared the amounts of sequestered cardenolides across treatments and milkweed bug species by ANOVA followed by LSMeans Tukey HSD test and included bug species and treatment as model effects. Additionally, we estimated Pearson’s correlation coefficients between body mass and concentration of sequestered cardenolides in *O. fasciatus*. Body length data were analysed by ANOVA followed by LSMeans Tukey HSD test, including sex, treatment, and the interaction between sex and treatment as model effects. Lifespan and number of hatchlings were analysed by ANOVA followed by LSMeans Differences Student’s t-test, including treatment as the model effect. For *O. fasciatus*, ‘experiment’ was always included as a model effect in our statistical analysis. Sample sizes for every experiment are mentioned in the figure legends. Probability values < 0.05 were considered statistically significant.

## 3. Results

### 3.1. Influence of dietary cardenolides on growth

We examined the influence of dietary toxins on the growth of *P. apterus, O. fasciatus, C. nerii, S. pandurus*, and *A. longiceps* (Figure 2) using an artificial diet containing increasing doses of cardenolides (Supplementary Figure 1). Growth of *P. apterus* was compromised substantially [F(3, 36) = 8.83, P < 0.001] upon exposure to dietary cardenolides (P < 0.001, LSMeans Tukey HSD). In contrast, *S. pandurus* [F(3, 35.81) = 1.5, P = 0.23] and *A. longiceps* [F(3, 40) = 1.11, P = 0.36] grew equally well across all diets. Remarkably, cardenolides had a positive effect on growth in *O. fasciatus* [F(3, 88.01) = 5.33, P = 0.002] and *C. nerii* [F(3, 36) = 5.69, P = 0.003]. We observed increased growth in the presence of dietary toxins across all doses in *O. fasciatus* (low vs. control, P = 0.008; medium vs. control, P = 0.005; high vs. control, P = 0.015, LSMeans Tukey HSD). Compared to a diet without cardenolides, *C. nerii* grew better on the low (P = 0.002) and the high-dose (P = 0.02), but not on the medium-dose diet (P = 0.09). Since our lab strain of *O. fasciatus* was highly inbred, we carried out the same experiment with a different lab strain of *O. fasciatus* and obtained similar results (Supplementary Figure 3). The amount of excretion products was not influenced by the presence of toxins across diets all bug species, *O. fasciatus* [F(3, 19) = 8.34, P < 0.001], *C. nerii* [F(3, 14) = 1.38, P = 0.29], *S. pandurus* [F(3, 15) = 0.53, P = 0.67], and *A. longiceps* [F(3, 5) = 0.96, P = 0.48], except *P. apterus* (lower in the presence of cardenolides, F(3, 15) = 4.69, P = 0.02) suggesting that stronger growth in *O. fasciatus* and *C. nerii* is not due to increased food uptake mediated by a phagostimulatory effect of dietary cardenolides (Supplementary Figure 4).

**Figure 2.**
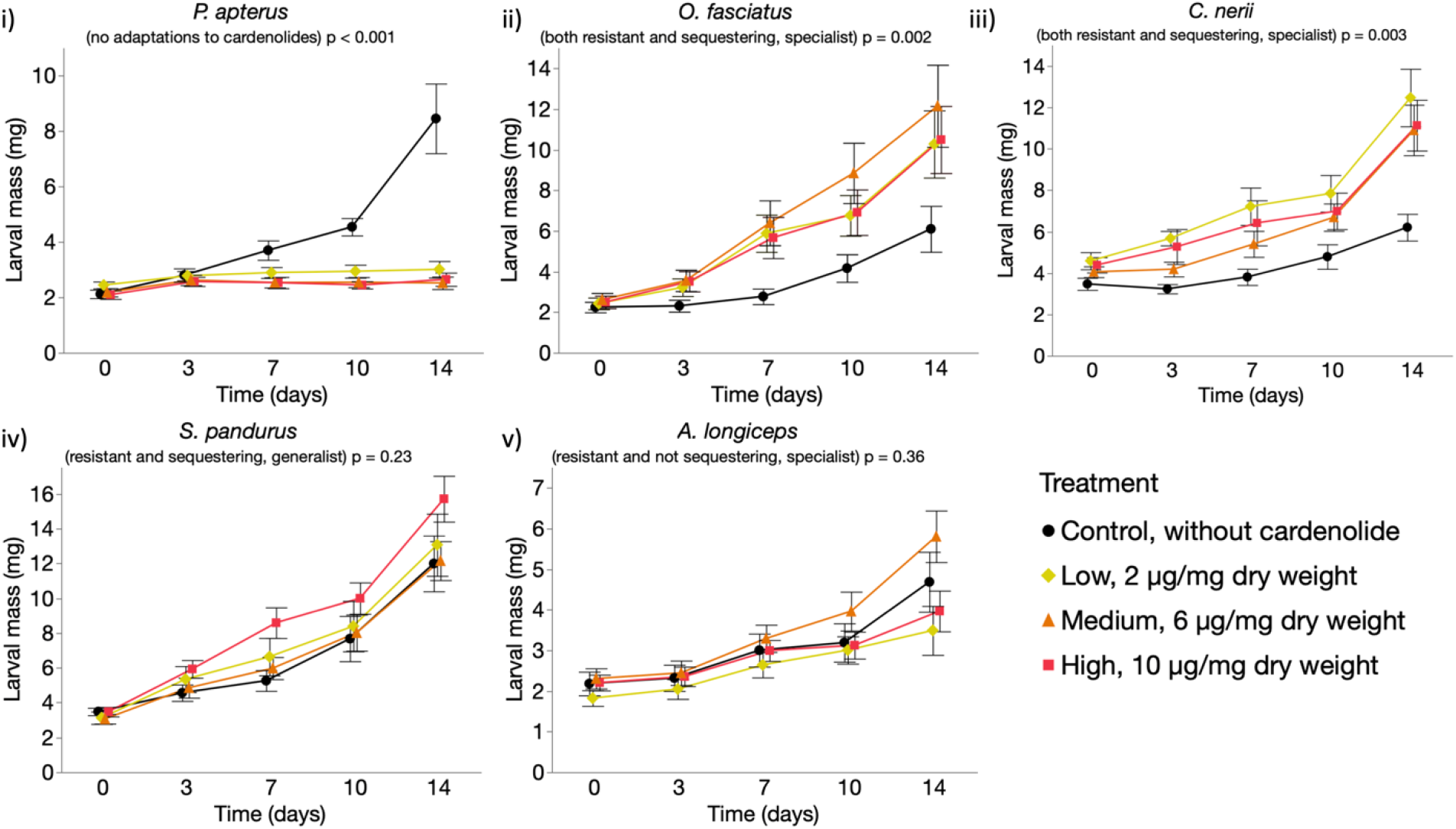
Growth of bugs on artificial diet with increasing doses of cardenolides. Each data point represents the mean (± SE) of larval mass at a given time. i) *P. apterus* (n = 10 per treatment), ii) *O. fasciatus* (n = 25 per treatment, three batches), iii) *C. nerii* (n = 10 per treatment), iv) *S. pandurus* (n = 10 per treatment), and v) *A. longiceps* (n = 10-13 per treatment).

### 3.2. Cardenolide sequestration

*P.apterus* did not sequester any cardenolides (Figure 3 i). We found substantial differences regarding the concentration of sequestered cardenolides across all treatments for *S. pandurus* [F(2, 12) = 12.11, P < 0.001], *O. fasciatus* [F(2, 27) = 15.99, P < 0.001], and *C. nerii* [F(2, 12) = 18.16, P < 0.001]. Among all the bug species, *O. fasciatus* sequestered remarkably higher amounts of cardenolides (P < 0.001). Although *A. longiceps* possesses a resistant Na^+^/K^+^-ATPase, we observed only very small concentrations of sequestered cardenolides which is consistent with earlier findings (Bramer et al. 2015). *S. pandurus* sequestered similar amounts of cardenolides from the diet with the intermediate and with the highest concentration of cardenolides (P = 0.89). We found dose-dependent cardenolide sequestration in *O. fasciatus* (low-medium, P = 0.011; low-high, P < 0.001; medium-high, P = 0.05) and in *C. nerii* (low-medium, P = 0.01; low-high, P < 0.001; medium-high, P = 0.05). Additionally, the sequestration data in *O. fasciatus* revealed an inverse relationship between body mass and concentration of sequestered cardenolides [low, r (10) = −0.7, P = 0.03; medium, r (10) = −0.92, P < 0.001; high, r (10) = −0.98, P < 0.001] (Figure 3 ii).

**Figure 3.**
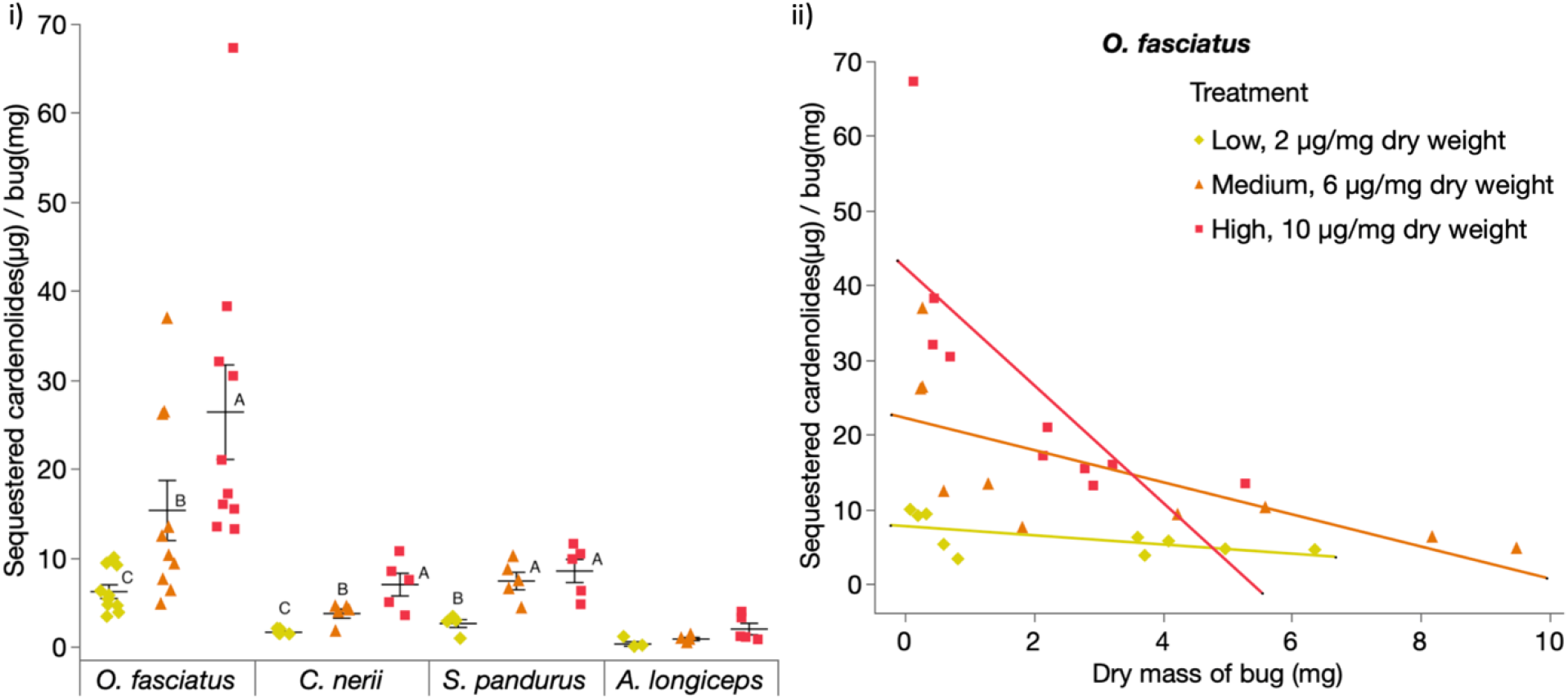
Sequestration of cardenolides by four species of milkweed bugs on artificial diet with increasing amounts of cardenolides. i) Total amounts of cardenolides sequestered per species [n = 5, per treatment for all species except of *O. fasciatus* (n = 10)] and dietary cardenolide concentration. Horizontal bars represent the mean concentration (± SE) of sequestered cardenolides. Within the same bug species, different letters indicate significant differences across treatments and dots represent jittered raw data. ii) Correlations between dry body mass and concentration of sequestered cardenolides in *O. fasciatus* (n = 10 per treatment). Trend lines represent the linear least squares regression fits to data points and dots represent jittered raw data.

In addition to the total amounts of cardenolides sequestered, we also compared the number of structurally different cardenolides (i.e. putative cardenolide metabolites) across the sequestering species (Supplementary Figure 2). Based on retention time comparison using an authentic standard, ouabain was sequestered as such. However, a peak with the retention time of digitoxin was not detected, but we found up to three compounds with a cardenolide spectrum and increased polarity probably representing digitoxin metabolites. Remarkably, we found more than one and up to three putative digitoxin-metabolites in *S. pandurus* and *C. nerii*. In *O. fasciatus*, we did not find any digitoxin-metabolites.

### 3.3. Influence of dietary cardenolides on lifespan, longevity and body size of *O. fasciatus*

Dietary cardenolides showed substantial effects on the developmental speed and longevity of *O. fasciatus*. Larvae raised on cardenolide-containing diet developed faster into adults [F(1, 26) = 42.79, P < 0.001, LSMeans Differences Student’s t test] and adults resulting from larvae raised on cardenolide-containing diet lived longer [F(1, 26) = 6.39, P = 0.02, LSMeans Differences Student’s t test] compared to individuals raised on cardenolide-free diet (Figure 4 i). Furthermore, cardenolides affected adult body size [F(3, 36) = 22.31, P < 0.001]. Females raised on toxic diet were the largest across all combinations (i.e. sex vs. diet; P < 0.001, when compared to males on toxic or control diets and P = 0.02, when compared to females on the control diet, LSMeans Tukey HSD). The body length of male *O. fasciatus* was not different between the two diets (P = 0.74, LSMeans Tukey HSD) (Figure 4 ii).

**Figure 4.**
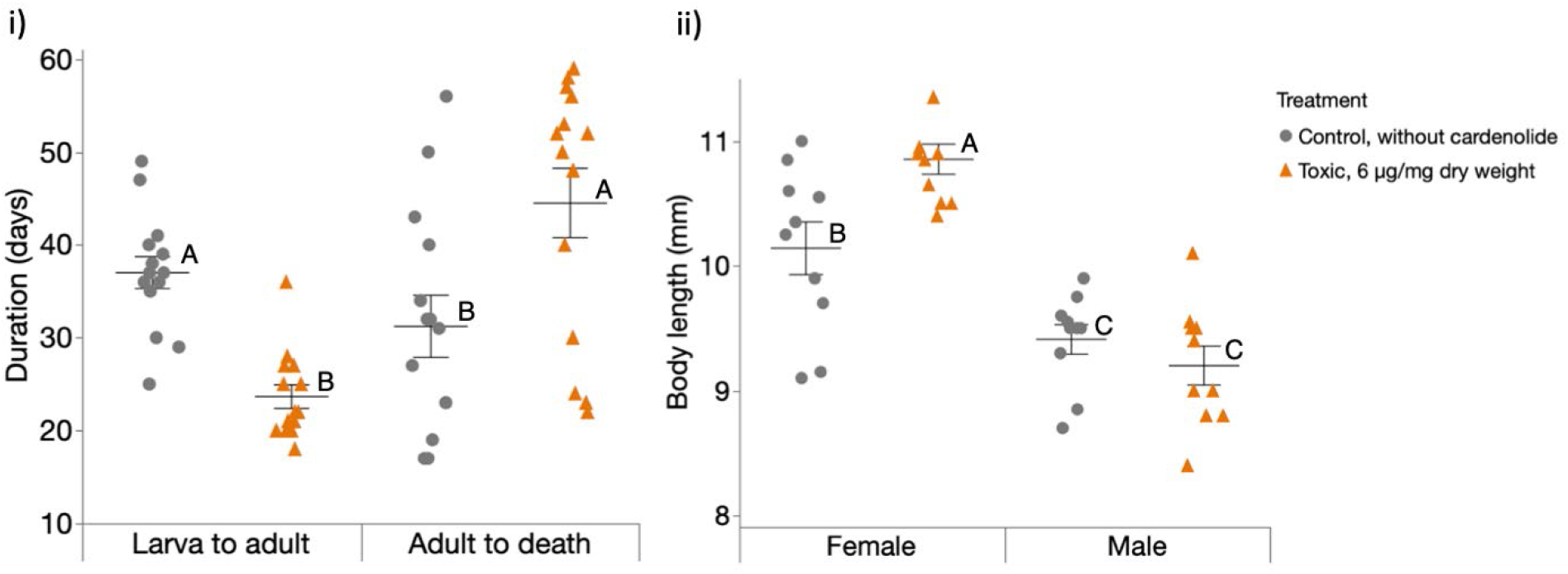
Effect of cardenolides on developmental time, lifespan and body size of *O. fasciatus*. i) Each horizontal line represents the mean (± SE) number of days taken by *O. fasciatus* larvae (n = 20 per treatment) to turn adult (left side), and until death after reaching adulthood (right side); within the same category (left or right), different letters indicate significant differences between diet treatments. ii) Each horizontal line represents the mean (± SE) body length of adult (female or male) *O. fasciatus* (n = 10 per treatment); different letters above bars indicate significant differences. Larvae were either raised on toxic diet or a control diet. Dots represent jittered raw data.

### 3.4. Influence of dietary cardenolides on reproductive fitness of *O. fasciatus*

We observed substantial effects on the reproductive fitness of female *O. fasciatus* upon exposure to dietary cardenolides. When we continued feeding females after reaching adulthood on the same diet they fed upon as larvae, *O. fasciatus* on cardenolide-containing diets produced less hatchlings than individuals raised on cardenolide-free diet [F(1, 23) = 15.82, P = 0.001, LSMeans Differences Student’s t test]. In contrast, *O. fasciatus* from cardenolide-containing and cardenolide-free diets produced similar numbers of hatchlings, when both groups were fed with sunflower seeds after reaching adulthood [F(1, 21) = 0.52, P = 0.48, LSMeans Differences Student’s t test] (Figure 5).

**Figure 5.**
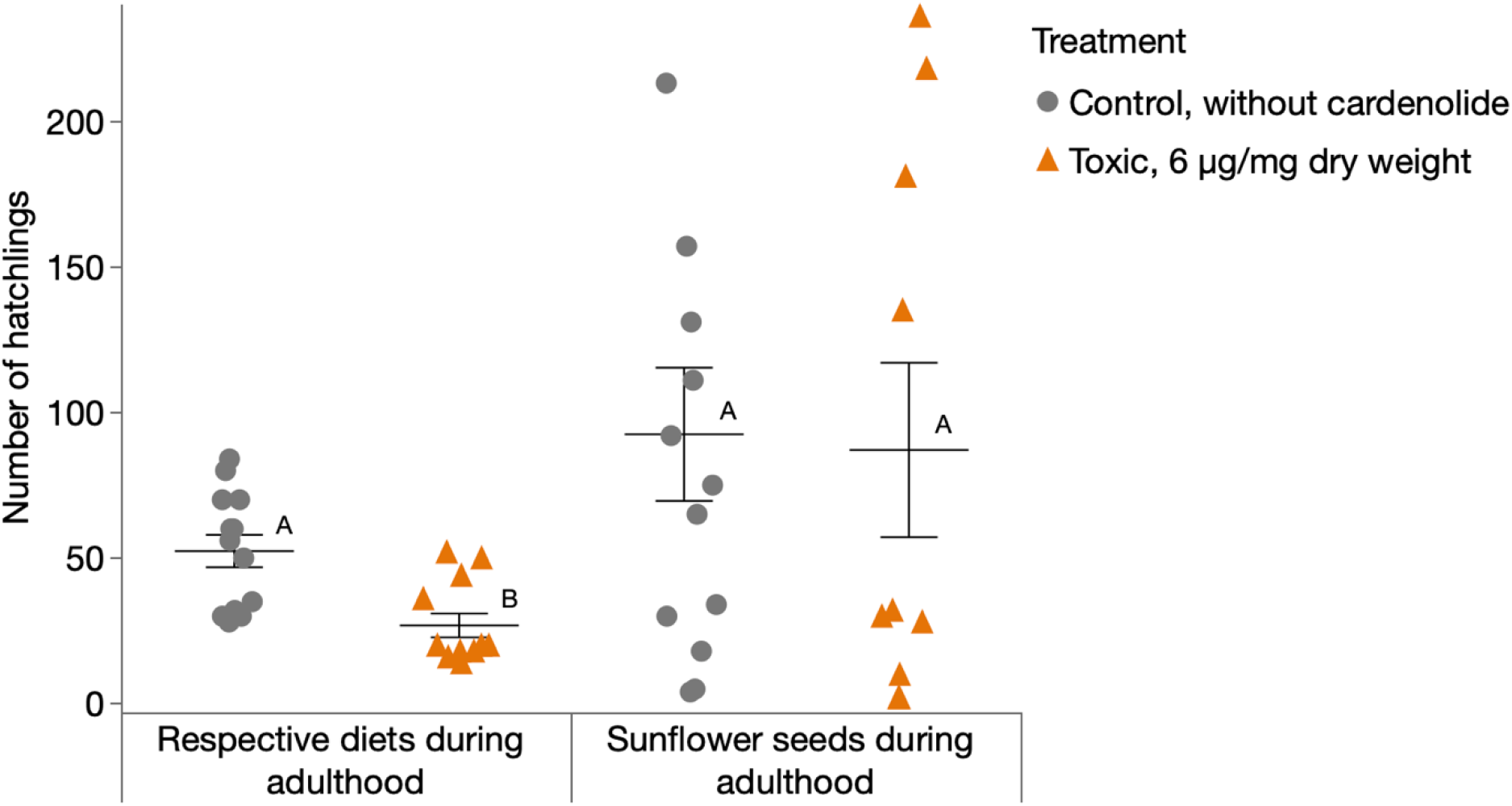
Reproductive success of *O. fasciatus* in the presence or absence of dietary cardenolides. Each horizontal bar represents the mean (± SE) total number of hatchlings produced by *O. fasciatus* females until death. Larvae were either raised on toxic diet or control diet until adulthood. A group of bugs stayed on the respective diets after reaching adulthood (left, n = 15 pairs per treatment) while the other group was transferred to sunflower seeds (right, n = 25 pairs per treatment). Within the same group, different letters indicate significant differences. Dots and triangles represent jittered raw data.

## 4. Discussion

We investigated if dietary cardenolides affected the growth of closely related species of milkweed bugs (*O. fasciatus*, *C. nerii*, *S pandurus*, and *A. longiceps*) and the outgroup species, *P. apterus* possessing different combinations of the traits ‘cardenolide resistance’ and ‘cardenolide sequestration’ and having different dietary strategies (generalist vs. specialist). Remarkably, dietary cardenolides increased growth in the sequestering specialists *O. fasciatus* and *C. nerii*, but not in the sequestering generalist *S. pandurus*. *O fasciatus* nymphs completed their development two weeks earlier and lived on average 13 days longer than adults under cardenolide exposure but produced less offspring unless being transferred to a cardenolide-free diet (sunflower seeds) after reaching adulthood.

Empirical evidence on the effect of dietary plant toxins on insect growth is ambiguous as different studies suggest contradictory effects. Nevertheless, several studies found effects that are in agreement with our study. For example, the growth of *O. fasciatus* was faster when raised on *Asclepias* species containing higher amounts of cardenolides (*A. syriaca* and *A. hirtella*) than when raised on species with low cardenolide contents (*A. incarnata* and *A. viridiflora*) (Chaplin and Chaplin 1981). Additionally, the African danaid butterfly *Danaus chrysippus*, having a cardenolide-resistant Na^+^/K^+^-ATPase (Petschenka et al. 2013), developed faster and produced larger adults when reared on *Calotropis procera* containing cardenolides compared to caterpillars raised upon *Tylophora* spp. lacking cardenolides (Rothschild et al. 1975). Furthermore, *Zygaena filipendulae* larvae reared on the plant *Lotus corniculatus* containing cyanogenic glucosides developed faster, and the larvae showed decelerated development when reared on transgenic *L. corniculatus* free of cyanogenic glucosides (Zagrobelny et al. 2007).

In contrast to these studies, caterpillar growth of the milkweed butterfly species *Euploe core*, *D. plexippus*, and *D. gilippus* was unaffected by cardenolides of eight *Asclepias* species ranging from very low to very high cardenolide contents (Petschenka and Agrawal 2015). Notably, all the studies mentioned above focused on insect feeding performance on intact plants or plant organs such as leaves or seeds that naturally represent highly complex diets. This could be one reason why contradicting outcomes in response to the same class of chemical compounds were observed even within the same plant species. Here, we used an artificial diet to control for variation across dietary treatments rigorously.

We showed that cardenolides had a positive effect on growth in *O. fasciatus* and *C. nerii*, both of which may be categorized as dietary specialists feeding on seeds of *Asclepias* spp. and seeds of *Nerium oleander*, respectively. We speculate that the positive impact on growth upon exposure of toxins may be due to selection of the resistant Na^+^/K^+^-ATPases to function optimally in a ‘toxic environment’, i.e. in the body tissues of a milkweed bug storing large amounts of cardenolides. In other words, there could be functional trade-offs of cardenolide-adapted Na^+^/K^+^-ATPases in a physiological environment that is devoid of cardenolides, a phenomenon that could be called ‘evolutionary addiction’. Alternatively, better growth could be due to increased consumption of the larvae mediated by cardenolides as phagostimulants (Pantle and Feir 1976). Nevertheless, our excretion data hint towards equal consumption of diet regardless of dietary cardenolide concentration. (Supplementary Figure 4).

The milkweed bug species *A. longiceps* is specialised on plants producing no cardenolides such as *Platanus* spp. or *Ulmus* spp. However, due to its evolutionary history, *A. longiceps* possesses resistant Na^+^/K^+^-ATPases, but has lost the ability to sequester cardenolides (Bramer et al. 2015). In contrast, *S. pandurus* possesses resistant Na^+^/K^+^-ATPases, and feeds on a wide array of host plants (Péricart 1998; Vivas 2012), including cardenolide producing species such as *Nerium oleander* and species of *Calotropis* from which it sequesters cardenolides (Von Euw, Reichstein, and Rothschild 1971; Abushama and Ahmed 1976). For both species, dietary cardenolides did not influence growth, i.e. they grew equally well on all diets.

The lack of a positive effect of dietary cardenolides on growth in *A. longiceps* may be associated with the inability of this species to sequester cardenolides. Accordingly, its Na^+^/K^+^-ATPases may have undergone a different selection regime compared to the sequestering species. In other words, their putative suite of Na^+^/K^+^-ATPases may have adapted to the absence of cardenolides in the body tissues secondarily. Alternatively, or in addition, there could be further physiological mechanisms involved mediating between sequestration and growth such as gaining energy from metabolising sequestered toxins. Nevertheless, the latter seems rather unlikely also as an explanation for increased growth in *O. fasciatus* and *C. nerii*, since the sugar moieties in digitoxin are dideoxy sugars not known to be easily metabolised as an energy resource (Liu and Thorson 1994).

While it is not surprising, that cardenolide-resistant Na^+^/K^+^-ATPases alleviate toxicity of dietary cardenolides in both species, *A. longiceps* and *S. pandurus*, it is an open question why the sequestering *S. pandurus* did not show increased growth under cardenolide exposure. Although *O. fasciatus* sequesters substantially higher amount of cardenolides in comparison to *S. pandurus*, it seems unlikely that the extent of sequestration is the underlying mechanism, given that *C. nerii* sequesters concentrations similar to *S. pandurus*. Moreover, for *O. fasciatus*, we showed that the amount of sequestered cardenolides is directly proportional to its body mass. Along the same lines, cardenolide metabolism may also not explain the patterns of growth since we found pronounced differences in metabolism between *O. fasciatus* and *C. nerii* both showing increased growth under cardenolide exposure. However, the putative adaptations underlying generalist feeding behaviour of *S. pandurus* likely caused different selection pressures in this species and thus interfere with the pattern of amino acid substitutions across the different Na^+^/K^+^-ATPases or lead to differential expression of the Na^+^/K^+^-ATPase genes across different tissues.

The outgroup species, *P. apterus* belongs to a different family (Pyrrhocoridae) and is not adapted to cardenolides, i.e., it has a cardenolide sensitive Na^+^/K^+^-ATPase and is not able to sequester cardenolides (Bramer et al. 2015). The lack of a resistant Na^+^/K^+^-ATPase most likely explains why larval growth in this species was compromised substantially by dietary cardenolides. Alternatively, or in addition, reduced growth could be due to feeding deterrence mediated by dietary cardenolides as indicated by the reduced amount of excretion products observed during our feeding trial.

Although specialist insects can successfully feed on toxic host plants, it is generally expected that the underlying resistance traits incur costs via trade-offs, which are expected to depend on the ecological context and the molecular mechanisms involved (Peterson, Hardy, and Normark 2016; Smilanich, Fincher, and Dyer 2016). As *O. fasciatus* raised on cardenolide-containing diet developed faster into adults and had a longer lifespan compared to those raised on cardenolide-free diet, cardenolide exposure clearly influences the fecundity-longevity trade-off. A faster development and an extended lifespan both indicate higher fitness. Nevertheless, individuals exposed to dietary cardenolides produced a lower number of hatchlings contradicting higher fitness. This apparent disadvantage, however, could be alleviated by maternal transfer of cardenolides to the eggs mediating protection against predators (Newcombe et al. 2013). Moreover, we found that feeding on a non-toxic diet (i.e. sunflower seeds) as adults can compensate for this effect. This likely resembles the natural situation since *O. fasciatus* larvae feed and cluster around on *Asclepias* seeds or seed-pods, while adults disperse and forage (Feir 1974). In conclusion, it seems likely that cardenolides exert a positive effect on overall fitness in *O. fasciatus* which is in disagreement with theory predicting costs of sequestration.

Dealing with xenobiotics can require energy allocation towards metabolism or can cause pleiotropic effects due to particular molecular mutations that might confer a selective advantage in the presence of xenobiotics but incur cost in their absence (Coustau, Chevillon, and ffrench-Constant 2000; Mauro and Ghalambor 2020). Although it is unclear how the function of Na^+^/K^+^-ATPase could be related to growth and longevity in milkweed bugs the enzyme is involved in many physiological processes apart from being a cation carrier suggesting many unknown non-canonical functions of the enzyme (Liang et al. 2007) which could provide a mechanistic link. In *O. fasciatus*, the three gene copies (α1A, α1B and α1C) encoding α-subunit of cardenolide resistant Na^+^/K^+^-ATPases have diverse functions (Lohr et al. 2017) and duplicated α1 gene copies do not only vary in number and identity, but also show specific expression patterns in different body tissues (Zhen et al. 2012; Yang et al. 2019; Dobler et al. 2019) allowing for complex regulation.

In conclusion, reduced growth in the absence of cardenolides in highly specialised milkweed bugs with cardenolide-resistant Na^+^/K^+^-ATPases suggests a novel type of physiological cost arising in the absence of plant toxins. Mechanistically, this cost could be due to negative pleiotropic effects mediated by resistant Na^+^/K^+^-ATPases not functioning optimally in a physiological environment lacking cardenolides. Furthermore, the observed effects of cardenolides on the fecundity-longevity trade-off probably leading to increased fitness in *O. fasciatus* may be due to optimized resource allocation under the influence of sequestered cardenolides. Our study suggests a concept that is contradictory to the general assumption that sequestered plant toxins produce physiological costs and our results indicate a further level of coevolutionary escalation.

## Acknowledgements

We thank Sabrina Stiehler for technical support and Martin Kaltenpoth for his advice on rearing *Pyrrhocoris apterus*. We are greatly indebted to Andreas Vilcinskas and the Justus Liebig University Giessen for infrastructural support. This research was funded by DFG grant PE 2059/3-1 to G.P. and the LOEWE Program of the State of Hesse by funding the LOEWE Center for Insect Biotechnology and Bioresources.

## Author contributions

PP and GP designed the experiments; PP collected and analysed the data; PP and GP wrote the manuscript; AS contributed critically to the drafts. All authors gave final approval for publication.

## Supplementary Material

### Supplementary Method

#### Quantification of excretion products

We estimated the amount of food uptake by quantifying the amount of excretion products during our feeding assay. Specifically, the area of faecal stains on filter papers lining the Petri-dishes were analysed. We only analysed filter papers from Petri-dishes in which all three bugs survived until the end of the experiment (i.e. for three weeks). Filter papers were scanned and the stained area was quantified by following the instruction of image analysis (Reinking 2007) using ImageJ 1.52k (National Institutes of Health, US). Excretion data were log_10_-transformed to achieve homogeneity of variances and normality of residuals. To test for differences in excretion across treatments data were analysed by ANOVA followed by the LSMeans Tukey HSD test in JMP.

### Supplementary Results

#### Estimation of excretion area

We estimated the area of excretion products on filter paper to assess bug feeding activity during our feeding trials. In line with increased growth of *P. apterus* on the control diet [F(3, 15) = 4.69, P = 0.02], we observed statistical differences of excretion products. Specifically, bugs excreted less waste products when fed on medium (P = 0.01) and high-dose (P = 0.04) but not on low dose diet (P = 0.22) compared to the control. In contrast, we found no statistical differences of excreted waste products across treatments in *C. nerii* [F(3, 14) = 1.38, P = 0.29], *S. pandurus* [F(3, 15) = 0.53, P = 0.67], and *A. longiceps* [F(3, 5) = 0.96, P = 0.48] (Supplementary Figure 4). *O. fasciatus* [F(3, 19) = 8.34, P < 0.001] excreted similar amounts when fed on either control, low or medium-dose diet. Interestingly, *O. fasciatus* excreted less in the high-dose (but similar to control) than compared to low and medium-dose. Moreover, there was a difference between excretion on medium (P = 0.002) and low dose (P = 0.004) diet as compared to the high dose diet.

### Supplementary figures and legends

**Supplementary Figure 1.**
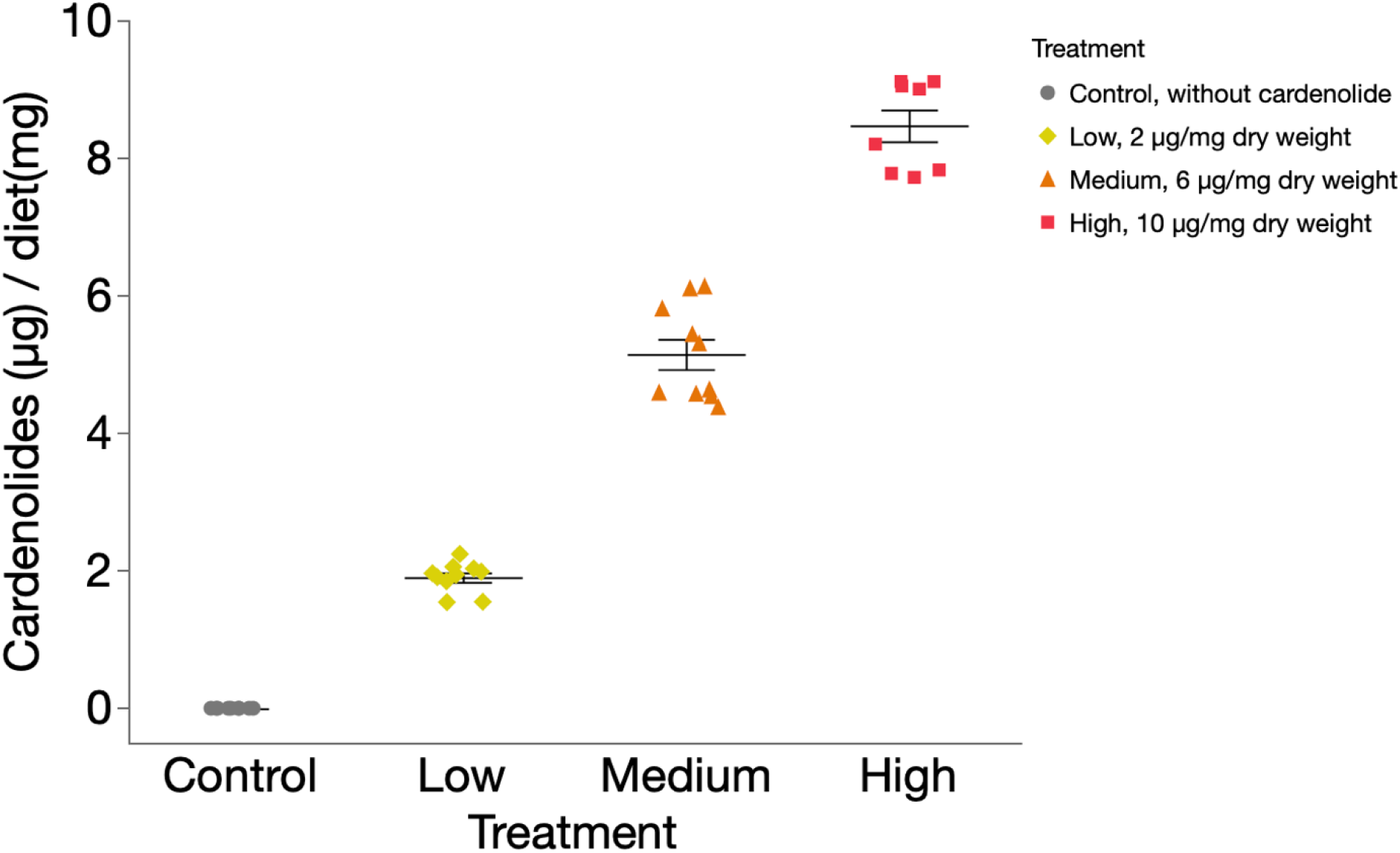
Concentration of cardenolides in the artificial diet. Each horizontal bar represents the mean concentration of cardenolides (±SE) in the diet (n = 10 per treatment). Symbols represent jittered raw data.

**Supplementary Figure 2.**
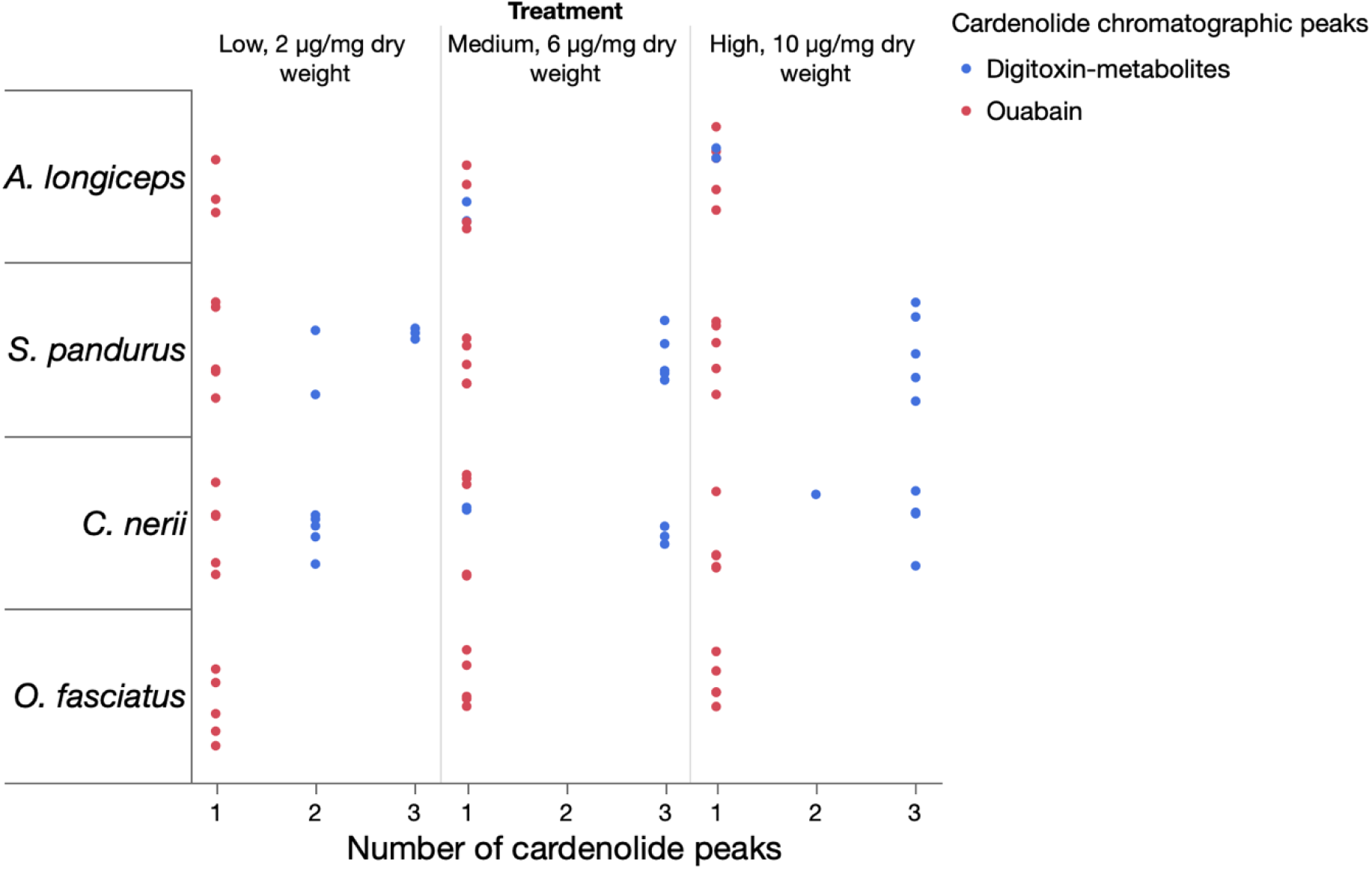
Structural diversity of sequestered cardenolide peaks by milkweed bugs. Each data point represents a bug specimen and the number of cardenolide peaks found in specimens of *O. fasciatus* (n = 10 per treatment), *C. nerii* (n = 5 per treatment), *S. pandurus* (n = 5 per treatment), and *A. longiceps* (n = 5 per treatment) when raised on artificial diet containing an equimolar mixture of ouabain and digitoxin. Chromatographic peaks with a cardenolide spectrum and a similar retention time like digitoxin were classified as digitoxin metabolites. Although we used a different HPLC method for *O. fasciatus*, the outcome is probably the same as if we had used the HPLC method used for *C. nerii*, *S. pandurus*, and *A. longiceps*. This assumption was validated by comparisons with *O. fasciatus* samples analysed during a different set of experiments (Heyworth et al., manuscript in preparation), hence it is valid to compare *O. fasciatus* to other species.

**Supplementary Figure 3.**
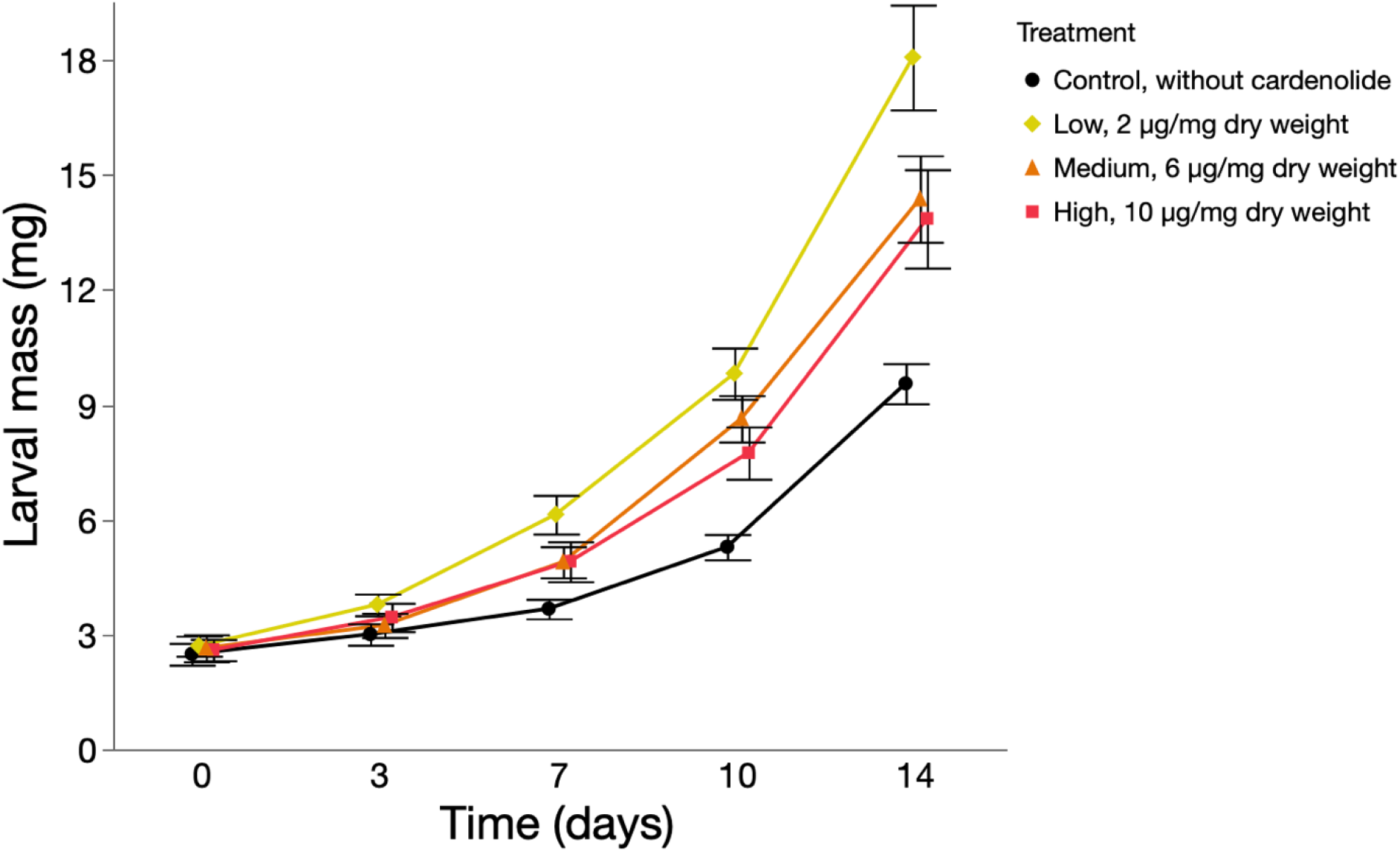
Growth of *O. fasciatus* on artificial diet with increasing doses of cardenolides using a genetically distinct *O. fasciatus* lab strain. Each data point represents the mean mass (± SE) of larvae raised on an equimolar mixture of ouabain and digitoxin (n = 10-12 per treatment). We commercially obtained *O. fasciatus* eggs from Carolina Biological Supply Company (Burlington, NC, US). Larvae were maintained on sunflower seeds as described above and used only in this feeding assay. Overall, cardenolides had a positive effect on growth [F(3, 39) = 5.53, P = 0.003, Repeated Measures ANOVA], but only low-dose (P = 0.002) was statistically significant from control, and not the medium (P = 0.07) and high-dose (P = 0.13, LSMeans Tukey HSD).

**Supplementary Figure 4.**
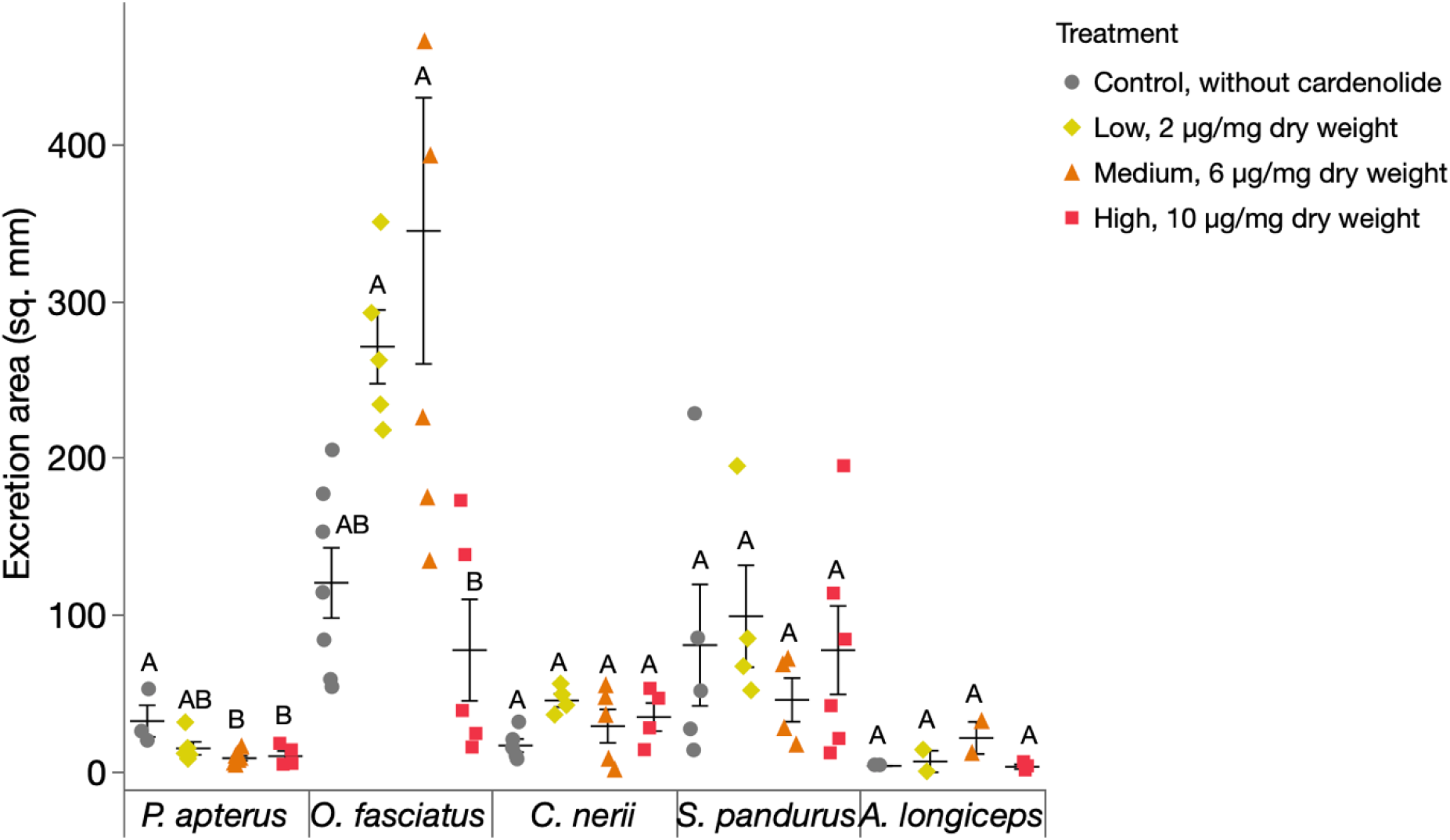
Amount of excretion products by bug species raised on increasing doses of cardenolides. Each horizontal bar represents the mean (± SE) of faecal stains on filter paper produced by *P. apterus* (n = 4-7 per treatment), *O. fasciatus* (n = 5-7 per treatment), *C. nerii* (n = 4-5 per treatment), *S. pandurus* (n = 5-6 per treatment), and *A. longiceps* (n = 2-3 per treatment) when raised on an equimolar mixture of ouabain and digitoxin. Within the same bug species, different letters indicate significant differences. Symbols represent jittered raw data.

